# *In situ* measurement of intracellular thermal conductivity using heater-thermometer hybrid diamond nanosensor

**DOI:** 10.1101/2020.06.03.126789

**Authors:** Shingo Sotoma, Chongxia Zhong, James Chen Yong Kah, Hayato Yamashita, Taras Plakhotnik, Yoshie Harada, Madoka Suzuki

## Abstract

Understanding heat dissipation processes at nanoscale during cellular thermogenesis is essential to clarify the relationships between the heat and biological processes in cells and organisms. A key parameter determining the heat flux inside a cell is the local thermal conductivity, a factor poorly investigated both experimentally and theoretically. Here, using a nanoheater/nanothermometer hybrid based on a polydopamine shell encapsulating a fluorescent diamond nanocrystal, we measured the intracellular thermal conductivity of HeLa cell with a spatial resolution of about 200 nm. Its mean value of 0.11 Wm^-1^K^-1^ determined for the first time is significantly smaller than that of water. Bayesian analysis of the data strongly supports the existence of variation of the intracellular thermal conductivity of about 40%. These results present a major milestone towards understanding the intracellular heat transfer phenomena at nanoscale.

## Introduction

Internal heat is a distinctive feature of mammals and birds^1–3^, but also occurs in plants^4–8^, insects^9^, fishes^10^ and possibly in dinosaurs^11^. The heat originates from intracellular biochemical sources and then flows through cells to elevate temperature of the whole body. The body temperature, in turn, is a factor governing enzymatic activities including those of heat sources^12^. Thus, simultaneous observation of biochemical processes at the heat source and the dynamics of heat flow *in situ* is essential for understanding the heat management in living organisms.

Luminescence nanothermometry enables visualization of temperature changes in cells with sub-cellular resolution^13,14^ and is a tool suitable for achieving this ambitious goal. Several groups used this tool and have recently reported temperature heterogeneity over 1 K in individual cells^15–18^. These reports have prompted active discussions about the origin and the biological significance of the heterogeneity. However, the results have also sparked hot debates about the validity of the temperature measurements^19–24^. The issue is that the temperature rise calculated by theoretical models describing physics of cellular thermogenesis and intracellular space is several orders of magnitude smaller than the experimental values reported in the literature. One of the essential factors in the calculation is the thermal conductivity, a parameter determining the heat flux and usually assumed to be about 1 Wm^-1^K^-1^, similar to a watery environment. However, Bastos *et al*. have used upconverting nanoparticles^25^ to measure the thermal conductivity of a single lipid bilayer and reported a value of 0.2 Wm^-1^K^-1^. Theoretically, the thermal conductivity in cells can be about six times smaller than that in water when the boundary heat resistance, or Kapitza resistance is considered^24^. Although it is pivotal to obtain the thermal conductivity in living cells and assess its variation at nanoscale experimentally, no such information is available to the best of our knowledge.

A two-in-one nanodevice which provides heating and temperature sensing at the same time and the same spot is a straightforward strategy for measuring intracellular thermal conductivity locally. Tsai *et al*. has reported the use of fluorescent nanodiamond (FND) and gold nanorods (GNRs)^26,27^. FNDs containing negatively charged nitrogen vacancy centers (NVCs) are known as a fluorescent nanomaterial with exceptional photostability, showing neither photobleaching nor photoblinking^28,29^. In addition, their unique magneto-optical properties allow nanoscale temperature measurement through optically detected magnetic resonance (ODMR) technique with accuracy better than 1 K^30,31^. Since NVCs are deeply embedded inside a diamond crystal with high thermal conductivity (larger than 50 Wm^-1^K^-1^)^32^, its thermosensing ability is little influenced by environmental factors^33^. Physically bound FND and GNR serve as a nanoheater and nanothermometer at the same time^26,27^ because GNRs release heat when exposed to light via strong surface plasmon resonance^34,35^. However, the stoichiometry of FND/GNR ratio is difficult to control and GNR can be deformed by heating, which makes quantitative and reproducible experiments difficult. More sophisticated hybrids need to be developed to overcome these difficulties.

Herein, we propose FND and polydopamine (PDA) nanohybrids (**Fig. 1A**). PDA was originally introduced as a multifunctional polymer^36^, inspired by the composition of adhesive proteins in mussels. Under basic conditions, dopamine molecules polymerize to form a PDA layer on the surface of inorganic materials. PDA demonstrates a photothermal conversion efficiency of 40%, which is much higher than 22% reported for gold nanorods^37^. Therefore, PDA-coated nanoparticles (e.g. gold, magnetic, and silica nanoparticles) have been utilized for multifunctional hyperthermia applications, especially for cancer therapy^38–42^. Several papers have already reported PDA-coated FNDs (PDA-FNDs) as a platform for chemical functionalization^43,44^. The structure of PDA-FND is highly uniform in a batch and the thickness of PDA layer can be easily controlled by changing the reaction conditions^44^. However, photothermal effect of PDA on FND has not been investigated despite the fact that the hybrid has immense potential for direct measurements of the thermal properties of living cells.

**Figure 1.**
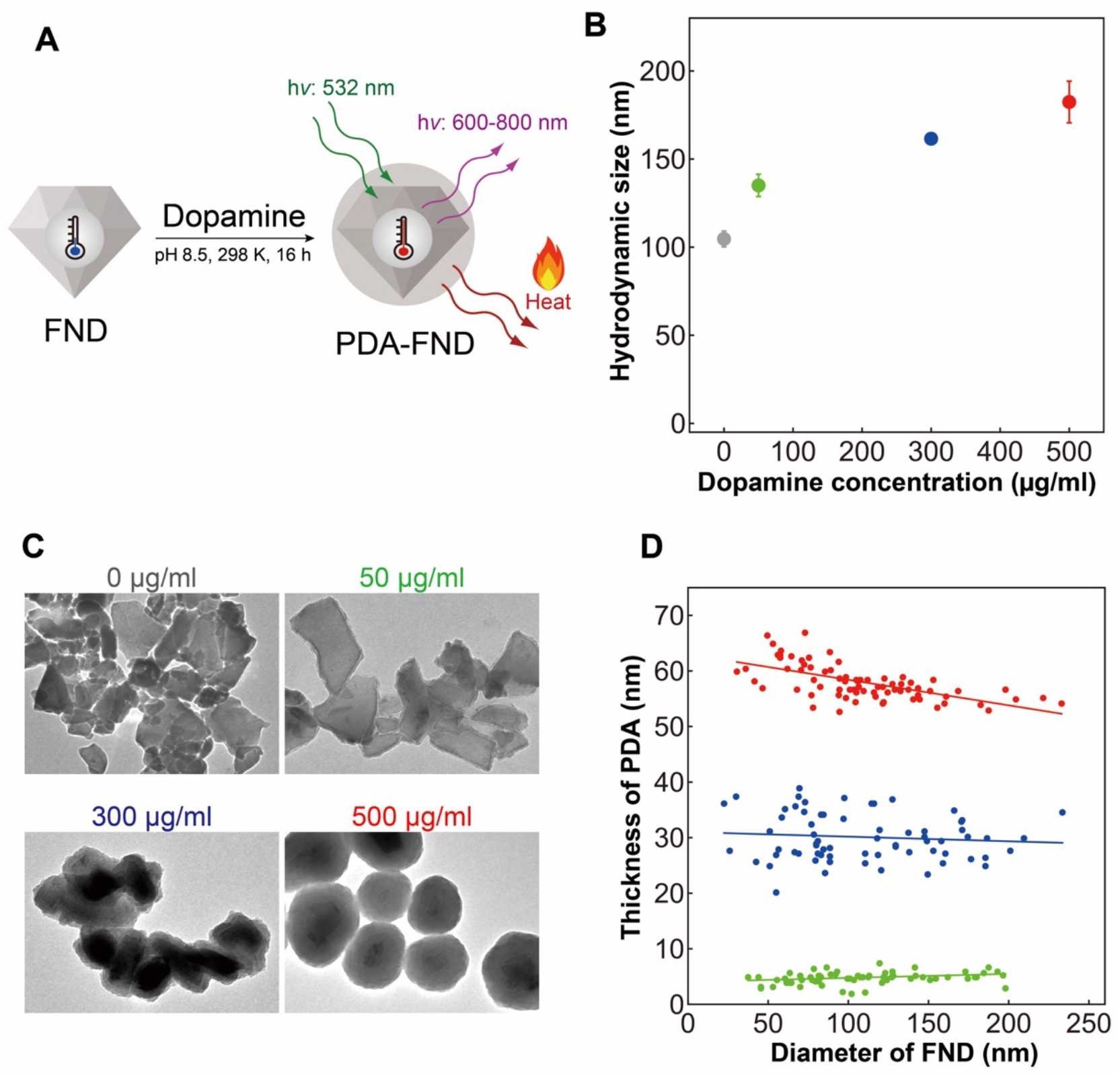
Characterization of PDA-FNDs. (A) Schematic illustration of the dual functionalized PDA-FND prepared from FND. The FND functions as a luminescent nanothermometer, while PDA releases heat by light-dependent manner. (B) Hydrodynamic diameters of FND and PDA-FNDs prepared with various concentrations of dopamine hydrochloride solutions. Plots and error bars indicate the average and the standard deviations (n = 5 independent preparations). (C) Representative TEM images of PDA-FNDs prepared with varied concentrations of dopamine hydrochloride solutions as indicated above each image. Dimension of images, 751 nm × 534 nm. Brightness and contrast were adjusted for viewing purposes. (D) Relationship between the diameter of FND and the thickness of PDA layer of PDA-FNDs prepared with 50 (green, n = 62 particles), 300 (blue, n = 68) or 500 (red, n = 75) μg/ml of dopamine hydrochloride solutions.

In this paper, we first characterize the physical properties of PDA-FND such as thickness of the PDA layer and the size of the FNDs and investigate how they affect the heat release. We confirm the structural stability of PDA-FND and reproducible heating cycles while measuring the temperature with accuracy around 1.0 K using individual PDA-FND particles. Then we validate our method by comparing the results of numerical simulations and the measured temperature increase when the particles dispersed on a substrate are in contact with air, water or oil for which the thermal conductivity coefficients are known from the literature. Finally, we measure thermal conductivity inside living single cells with a submicron spatial resolution.

## Results

### Synthesis and characterization of PDA-FNDs

Dopamine molecules are polymerized into PDA on the surface of FNDs in Tris-HCl buffer (pH 8.5), thus yielding PDA-FND after centrifugation and washing (**Fig. 1A**). The size, shape, and thickness of PDA layer of PDA-FNDs were evaluated by dynamic light scattering (DLS) and the transmission electron microscopy (TEM). The mean hydrodynamic diameter of bare FNDs was 105 nm, and PDA-FNDs prepared with 50, 300 and 500 μg/ml dopamine hydrochloride solutions had mean diameters of 135 nm, 162 nm and 182 nm, respectively (**Fig. 1B and Table S1**). TEM images show that FNDs have sharp edges and wide distribution in size and shapes (**Fig. 1C**). After PDA coating, FNDs have relatively smooth edges. Zeta potentials of the samples also support the successful synthesis of PDA-FNDs (**Fig. S1**). The thickness of PDA layer increases with the concentration of dopamine hydrochloride solution. The average thickness of the coating layer prepared with 50, 300, and 500 μg/ml of the dopamine hydrochloride solutions were 5 ± 1, 30 ± 4, and 58 ± 3 nm, respectively (**Fig. 1D**). The thickness of PDA layer appears to be constant regardless of the size of FNDs. The gradients of the straight lines in **Fig. 1D** are small (0.72%, - 0.83%, and −4.6% for 50-, 300- and 500-μg/ml samples, respectively).

### Dual functionality of PDA-FND nanoparticles

PDA-FND is a hybrid of luminescent nanothermometer and an optically controlled nanoheater. The ground state of NV^−^ center in FND is a spin-triplet. The split of the spin sublevels *m*_s_ = 0 and *m*_s_ = ±1 denoted as *D*_gs_ ≈ 2.87 GHz is temperature-dependent with a reported gradient^45^ *γ*≈-74 kHz K^−1^ (**Fig. S2**). Photoluminescence excited at 532-nm decreases when the transition from *m*_s_ = 0 to either of *m*_s_ = ±1 states occurs at resonance with applied external microwave (MW) radiation (**Fig. 2A**). The contrast of the ODMR signal decreases at larger excitation powers probably due to the photo-induced conversion of NV^−^ centers to ODMR-inactive neutral nitrogen-vacancy NV^0^ centers^46^.

**Figure 2.**
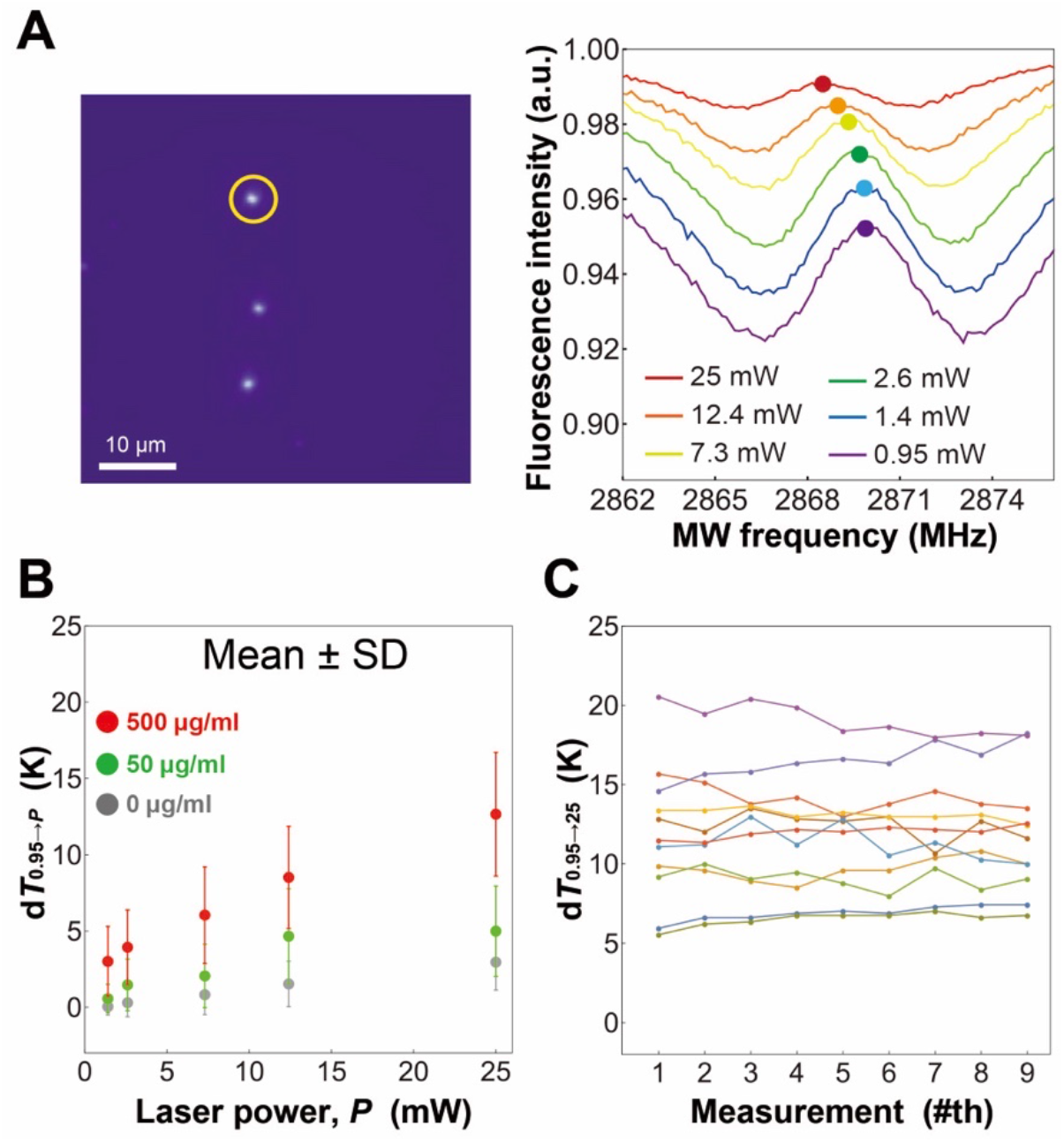
Application of PDA-FNDs as individual nanoheater/nanothermometer composites. PDA-FNDs for which data are shown in panels (A-C) are prepared using 500 μg/ml of dopamine chloride solution. (A) Typical fluorescence image of PDA-FNDs on a coverslip and ODMR spectra of the PDA-FND indicated by the yellow circle in the image recorded with different excitation laser powers. (B) Temperature change d*T*_0.95→*P*_ estimated from the change of *D*_gs_ in PDA-FND particles compared to pristine FNDs. The panel shows the corresponding mean values (circles) and the sample standard deviations (bars) which is presented in **Fig. S4**. (C) Values of d*T*_0.95→25_ (the temperature change when the laser power changes from 0.95 mW to 25 mW) in 11 individual PDA-FNDs obtained in 9 sequential measurements.

The values of *D*_gs_ vary from particle to particle in the range of 2869.0 – 2870.0 MHz at 0.95 mW excitation due to an inherent heterogeneity of FNDs^33,47^, but *D*_gs_ decreases at higher laser powers as expected in the case of rising temperature (**Fig S3**). Heterogeneity of *D*_gs_ prevents absolute measurements of temperature but one can obtain by monitoring the shift of *D*_gs_ values the temperature difference d*T*_*P*_1_→*P*_2__ caused by the change of the excitation power from *P*_1_ to *P*_2_. The values of d*T*_*P*_1_→*P*_2__ are about 4 times larger in particles prepared using 500 μg/ml dopamine hydrochloride solution in comparison to pristine FNDs (**Figs 2B** and **S4**). The PDA-FNDs prepared under 50 μg/ml dopamine hydrochloride solution showed intermediate photothermal efficacy, which means that the thin PDA can serve as a scaffold for further functionalization with some precaution related to heating. The result safeguards some applications using a thin layer of PDA against undesirable heating of the functionalized material^48–50^. Because PDA-FNDs prepared from 500 μg/ml solution have a large photothermal effect but do not significantly hinder FND fluorescence, they were used in subsequent experiments.

To assess the photostability of PDA, ODMR was measured repeatedly in the same set of individual PDA-FNDs prepared from 500 μg/ml solution in nine successive measurements of d*T*_0.95→25_. The temperature rise remains nearly constant in the series (**Fig. 2C, Fig. S5**). These results indicate that PDA layer in PDA-FND remains stable after multiple excitations, and the heat release is reproducible under the current experimental condition.

### Cellular uptake and intracellular localization of PDA-FNDs

The biocompatibility, cellular uptake, and intracellular localization of PDA-FNDs were examined in HeLa cells. It has been reported that both FNDs^51,52^ and PDA nanoparticles^53^ are biocompatible with various types of cells. Our test with CCK-8 assay indicates that PDA-FNDs barely affect the viability of HeLa cells. (**Fig. 3A**). The amount of PDA-FNDs used in the ODMR experiments in the following section was 1 μg/ml or less, 15% of the smallest concentration in **Fig. 3A**.

**Figure 3.**
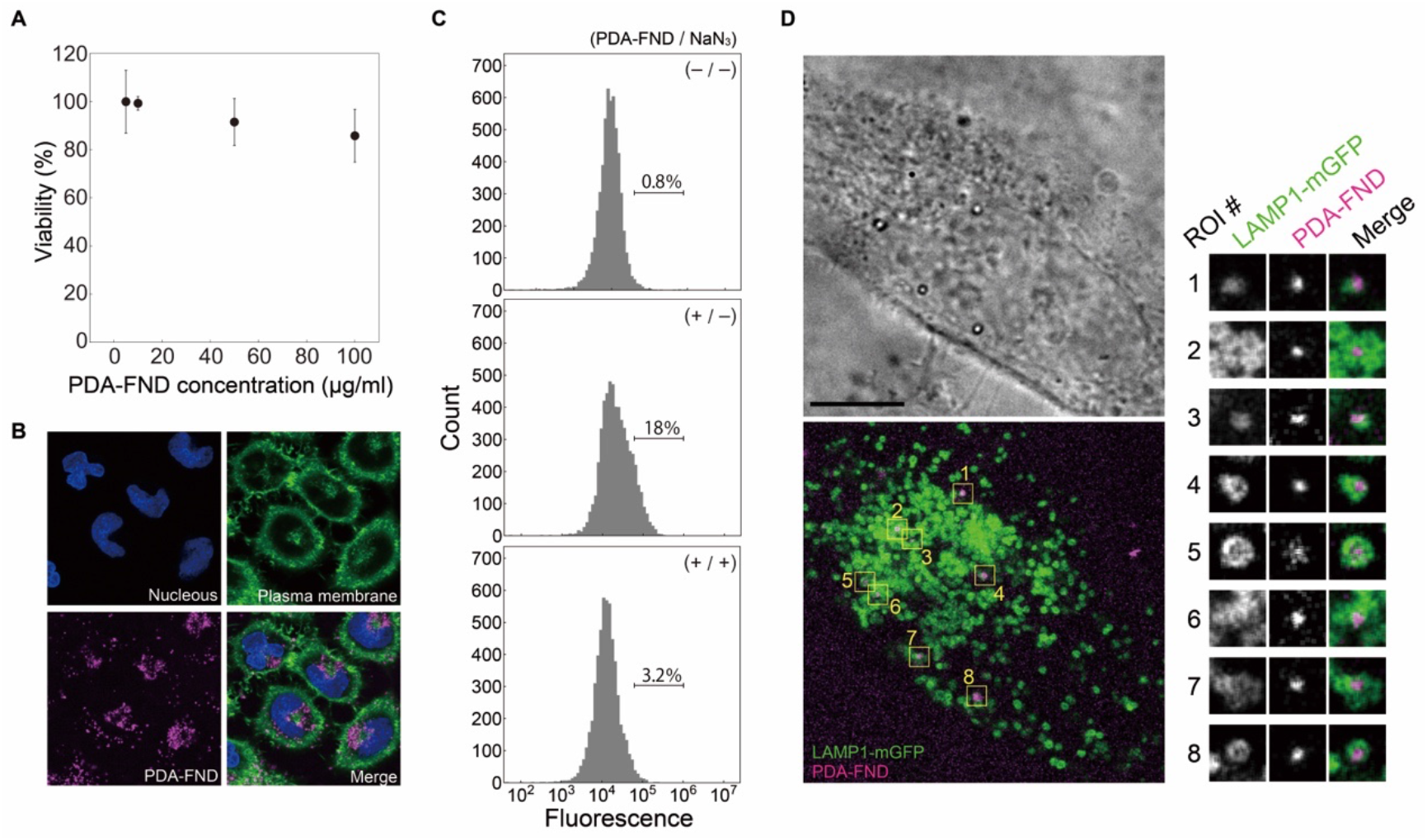
Biocompatibility and subcellular localization of PDA-FNDs. (A) Cell viability test of HeLa cells treated with PDA-FNDs at various concentrations using CCK-8 assay. (B) Confocal microscopic images of HeLa cells incubated with PDA-FNDs for four hours. Blue, nucleus stained by Hoechst 33342. Green, plasma membrane stained with CellMask Green. Magenta, PDA-FNDs. Maximum intensity of ten confocal sections at the middle of cells is overlaid. Images are smoothed, and the brightness and the contrast were adjusted for viewing purpose. Dimension of images; 92.3 μm × 92.3 μm. (C) FACS analysis of the cellular uptake of PDA-FNDs with and without NaN_3_ treatment. Non-treated HeLa cells (top), cells loaded with PDA-FNDs (middle), and cells loaded with PDA-FNDs in the presence of 10 mM NaN_3_ (bottom). The number on the bar represents the percentage of cells where the level of fluorescence is in the range from 5 × 10^4^ to 1 × 10^6^ (indicated by the bar). *Top*, the fraction of cells within the range indicates the number of cells where the level of autofluorescence is in the range. *Middle*, the fraction of cells within the range is elevated to 18%, implying the presence of cells containing PDA-FNDs. *Bottom*, the fraction of cells within the range is reduced to 3.1%. (D) Confocal microscopy images of HeLa cells overexpressing LAMP1-mGFP that were incubated with PDA-FNDs for four hours and chased for overnight. Left, bright field image showing a single optical section (top) and confocal fluorescence image showing the maximum intensity of six optical sections (bottom). Right, enlarged views of single optical sections from LAMP1-mGFP, PDA-FND and their merges. Areas correspond to the regions of interests (ROIs) shown in the image in center. All images are smoothed for viewing purposes. Scale bar, 10 μm. All the PDA-FNDs studied here were prepared with 500 μg/ml of dopamine hydrochloride solution.

PDA-FNDs were internalized by HeLa cells and they mainly localized at the perinuclear region after 4 hours incubation according to confocal observations. (**Fig. 3B**). FNDs are known to enter cells via ATP-dependent endocytosis, which is commonly observed among particles up to 150 nm in diameter^54^. In order to investigate the role of endocytosis in this case, we evaluated the effect of sodium azide (NaN_3_), an inhibitor of *F*_0_*F*_1_ ATP synthase, on the cellular uptake of PDA-FND. Confocal microscopy results showed that the amount of PDA-FNDs that was internalized by HeLa cells decreased remarkably in NaN_3_-treated cells compared to non-treated cells (**Fig. S6A**). Quantitative fluorescence activated cell sorting (FACS) analysis further supported the microscopic observations (**Fig. 3C**). The fraction of highly fluorescent cells was 18% after treatment with PDA-FND, which was markedly larger than that in control cells (0.8%). The fraction of cells containing PDA-FNDs was reduced to 3.2% when cells were treated with NaN_3_. These results strongly suggest that HeLa cells uptake PDA-FNDs through energy-dependent endocytosis.

Further investigation of the subcellular localization showed that most of PDA-FNDs are present in lysosome associated membrane protein 1 (LAMP1)-GFP positive-vesicles, i.e. in lysosomes, in LAMP1-mGFP[55] overexpressing cells (**Fig. 3D**). A small portion of PDA-FNDs in cells did not colocalize with LAMP1-GFP positive-vesicles and might either exist in endogenous LAMP1 vesicles or be distributed in cytosol (**Fig. S6B**). Because integrity of PDA layer is preserved under acidic condition in lysosome (**Fig. S7**), PDA-FNDs are suitable for intracellular measurements.

### *In situ* measurement of intracellular thermal conductivity

Temperature rise of a PDA-FND can be analytically found in the case of a spherical symmetry (**Fig. 4A**). The steady-state temperature *T*(*r*) follows the Fourier’s low;

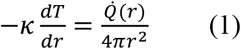

where *κ* is the thermal conductivity of the media. The heat power 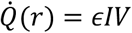, where *I* is the laser irradiance at the location of the particle, *ϵ* is the absorption coefficient of PDA, and *V* is the volume of PDA enclosed by radius *r* (dashed line in **Fig. 4A**). We will discuss the effect of the glass substrate later in this section. One can integrate **Eq. (1)** to obtain Δ*T*, the change of temperature in the FND caused by the change of irradiance Δ*I*. It reads;

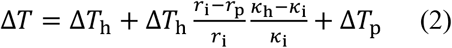

where

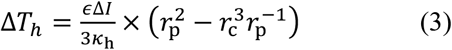

and

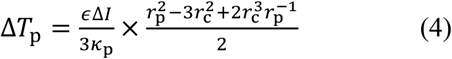

**Figure 4.**
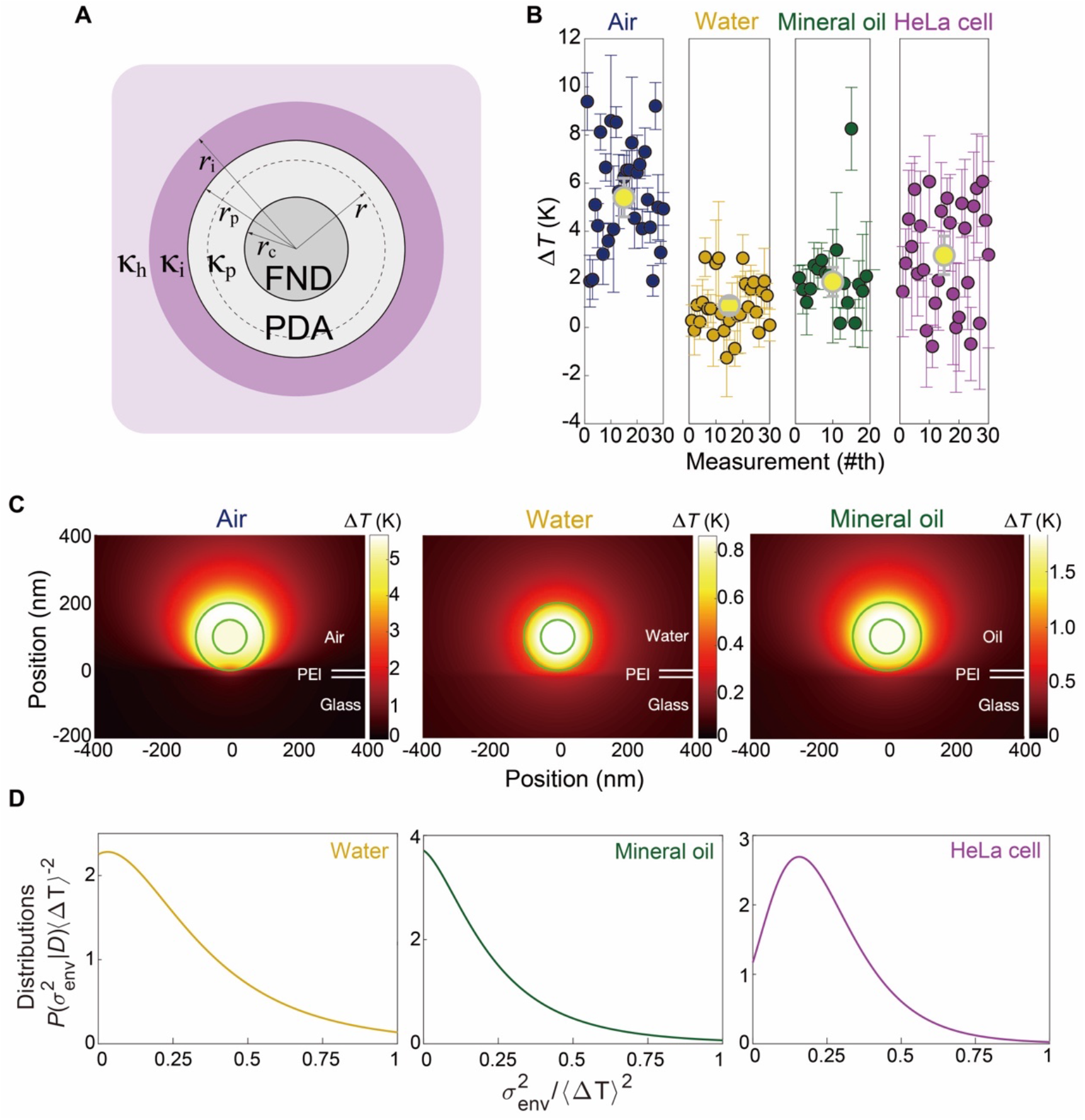
*In situ* measurement of local thermal conductivity in air, water, oil and HeLa cell. (A) Model structure of a PDA-FND and its surroundings. The radius of the FND is *r*_c_, and the radius of the PDA-FND is *r*_p_. The thermal conductivity *κ* of PDA is *κ*_p_, *κ* = *κ*_i_ in the region with variation *r*_p_ < *r* < *r*_i_, The rest of the space is homogeneous, *κ* = *κ*_h_. (B) Plots and error bars indicate Δ*T* and their standard deviations measured for each particle in different environments measured when the laser power increases from 7.3 mW to 25 mW. The yellow circles and grey bars show the estimated value of 〈Δ*T*〉 and their 95% confidence intervals. (C) Numerical simulations of the temperature distributions in and around a PDA-FND in air, water and oil (from left to right). The thermal conductivities of air, water and oil are set as 0.026, 0.601 and 0.135 Wm^-^K^-1^, respectively. The thermal conductivity and the heat power density of PDA are 0.2 Wm^-1^K^-1^ and 100 μW μm^-3^, respectively, in all cases. A 20-nm-thick layer of PEI (thermal conductivity set as 0.2 WmK^-1^) is placed on the top of the glass substrate. Circles show the border lines of the FND and of the PDA shell. The diamond temperatures are 5.49, 0.87 and 1.81 K above the base temperature. Note the difference in the color scale. (D) *Posterior* Bayesian inference *P* (*σ*^2^_env_|*D*) divided by 〈Δ*T*_h_〉^2^ ≈ 〈Δ*T*〉^2^ for water, oil and cells. The *prior P* (*σ*^2^_env_) is uniform.

Δ*T_h_* depends on the thermal conductivity of the media, whereas Δ*T*_p_ is independent of the surroundings but depends only on the intrinsic values of the PDA-FND (**Supplementary Information**).

Two factors *κ*_p_, the thermal conductivity of PDA and 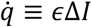, the power density of heat released by the PDA shell should be determined before using PDA-FNDs for measuring the unknown value, *κ*_h_. To do this, temperature rises were measured using different PDA-FNDs in air, water and oil as control media with known thermal conductivity values^56–58^. Bayesian inference is used to obtain *posterior* probability *P*(〈Δ*T*〉|*D*) (**Supplementary Information**, **Figs. S8-10**). In this case, the *data* (*D*) is the values of Δ*T* and their standard deviations measured for each particle (circles and error bars in **Fig. 4B**, respectively). ODMR spectra were obtained for PDA-FNDs attached to the bottom of the flow cell coated by polyethylenimine (PEI) with a thickness of 18.4±5 nm (95% confidence estimated using 22 measurements) (**Fig. S11**). The estimates obtained for d*T*_7.3→25_ which is denoted as 〈Δ*T*〉 in air, water and oil are shown in **Figs. 4B**, **S9** and given in **Table 1**. The two unknown parameters were tuned in numerical simulations to obtain 〈Δ*T*〉 which agrees with 〈Δ*T*〉 measured in all media simultaneously (**Fig. 4C**). Numerical simulations are required for accurate comparison between the data and the theory because the presence of the glass substrate (*κ*_glass_ ~ 1.05 Wm^-1^K^-1^)^59^ breaks the simplicity of the spherical symmetry considered in **Eq. (2)**, and the presence of PEI on the glass substrate cannot be neglected (**Table S2**). We have found that *κ*_p_ = 0.2 Wm^-1^K^-1^ and 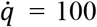 μW μm^-3^ (thus 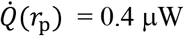) reproduce all values of 〈Δ*T*〉 (**Fig. 4C, Table 1**).

**Table 1.** 〈Δ*T*〉 obtained experimentally and by numerical simulations in different media with *κ*_p_, = 0.2 Wm^-1^K^-1^ and 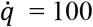 μW μm^-3^. A 20-nm-thick layer of PEI (*κ*_PEI_ = 0.2 Wm^-1^K^-1^) is placed on the top of the glass substrate; i.e. between the glass and the PDA-FND

| | $\langle \Delta T \rangle$ (experiment) | $\kappa$ (literature) | $\langle \Delta T \rangle$ (simulation) |
| --- | --- | --- | --- |
| | K | $\text{Wm}^{-1}\text{K}^{-1}$ | K |
| Air | $5.4 \pm 0.8$ | $0.026^a$ | 5.49 |
| Water | $0.9 \pm 0.3$ | $0.601^b$ | 0.87 |
| Oil | $1.9 \pm 0.6$ | $0.135^c$ | 1.81 |
| Cell | $3.0 \pm 0.8$ | $0.11^d$ | 3.01 |

<sup>a</sup> [Ref: 56]
<sup>b</sup> [Ref: 57]
<sup>c</sup> [Ref: 58]
<sup>d</sup> This work

Next, we measured temperature rises of individual PDA-FNDs inside cells. To avoid a cross-talk between PDA-FNDs, the concentration of PDA-FND solution applied to cells was reduced 20 times (to 0.5 μg/ml) compared to the amount used for studying intracellular localizations (cf. **Fig. 3D**). Very few or no PDA-FNDs were present in most of the cells. The mean value of temperature rise in cell 〈Δ*T*_cell_〉 was 3.0 K (**Fig. 4B**). Numerical simulations show that the influence of the glass substrate on the temperature of a PDA-FND is negligible when the distance between the probe and the glass is larger than 200 nm **(Fig. S12)**. In such a case, **Eq. (2)** is accurate. Therefore, *κ*_cell_ = 0.11 ± 0.3 Wm^-1^K^-1^ was calculated using **Eq. (2)** with *κ*_p_ and 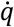 obtained above and the 〈Δ*T*_cell_〉. We note that 〈Δ*T*_cell_〉 in **Fig. 4B** visually spreads more than *(ΔT)* in other media. This observation has been examined quantitatively because it points to a significant variation of *κ*_cell_ due to complex intracellular environment.

The variance of Δ*T* calculated from **Eq. (2)** can be approximately split in two parts as follows (see **Eq. (S5)**).

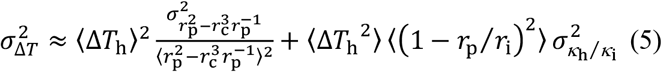

The first term in **Eq. (5)** is called 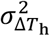 and is related to the statistics of PDA-FND nanoparticles independent of the surroundings. The second term, denoted 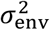 depends on the fluctuations of the local environment by means of *r*_p_/*r*_i_ and *κ*_h_/*κ*_i_. The method of obtaining the *posterior* distributions 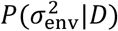 is explained in the Supplementary Information (**Eq. (S15)**). Distributions 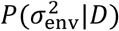 for water, oil and cell are shown in **Fig. 4D**. They are shown for 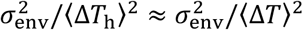 to eliminate the scaling effect of a larger temperature rise in media with smaller *κ*_h_. One can see that the peaks of 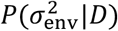 are near zero in water and oil as expected in media where it is reasonable to assume that 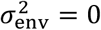. The result is a bit less conclusive for water (in oil 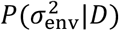 more sharply peaks at zero) where **Eq. (5)** is less accurate (see Supplementary Information). However, in cells, a clear peak is present at 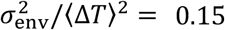. This indicates that fluctuations of *κ*_cell_ are significant. The variance of 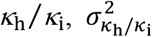, can be calculated using the value of 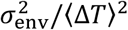 (**Eq. (S7)**). In the case *r*_i_ ≫ *r*_p_, one obtains 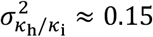. This value of the variance indicates that *κ*_cell_ can deviate from its average value by about 40% 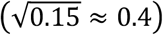.

## Discussion

The intracellular thermal conductivity *κ*_cell_, a key parameter characterizing the heat transfer in living cells has been experimentally determined in this paper. Single-cell measurement of *κ* has been previously attempted. Park *et al*. reported a value similar to that of water based on a three-omega method^60^, whereas the values of cancer cell lines determined by ElAfandy *et al*. using nanomembranes were slightly less than that of water^61^. We explain the difference by pointing out that both groups have provided heat to the specimen from outside of the cells. It is likely that the *κ* measured was an average of the specimen including a whole cell as well as extracellular matrix and nearby medium. In order to determine the intracellular *κ* at nanoscale, both heating and temperature measurement should be confined to a small volume inside cells, as we have demonstrated in the current study.

Bayesian analysis suggests that *κ*_cell_ is not a constant but can deviate from its average value. If one takes the 40% deviation into account, then the minimal value of *κ*_cell_ ≈ 0.07 W m^−1^K^−1^ and the steady-state Δ*T* ≈ 1 K (we set this value as a minimum temperature rise affecting biochemical processes) can be achieved in a spherical region generating 5 nW of heat power and approximately 10 nm across. While such a power is indeed produced by rat brown adipocyte^62^, the whole-cell power cannot be confined in a single hot spot. However, much less power is required to achieve a 1-K transient temperature rise. The heat conductivity, which is significantly smaller than the value previously assumed, reduces the speed with which the heat propagates through the intracellular medium. A transient value of Δ*T* in a region of radius *a* requires heat energy 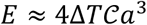, where 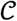 is the volumetric heat capacity. This temperature rise will last for a time interval 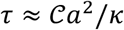. [24]. In the case 2*a* = 1 μm, Δ*T* = 1 K, 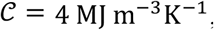, and *κ* = 0.07 W m^−1^K^−1^, the required energy is 2 pJ, about 10^-4^ times the energy of the glucose typically accumulated in a cell^19^ while the corresponding *τ* ≈ 15 μs. Notably, this time scale is characteristic for dynamics of some primary intracellular processes such as protein folding^63^. Therefore, the new value of *κ*_cell_ makes biologically significant transient temperature rises in a cell much more feasible. This conclusion suggests that cells may use heat flux for short distance thermal signalling.

Variation of thermal conductivity is expected to increase with decreasing the region where the thermal conductivity is measured. The intracellular environment has a complex architecture and composition. Various biomolecules at high-density form assemblies in the cytosol and organelles. In order to identify the source of the variation in *κ*_cell_, it is necessary to localise the accurate points where the temperature is measured. Technical challenges should be overcome to achieve this. One approach could include employing the catechol/quinone groups on the PDA as a versatile platform for functionalization^44,64,65^ to realize selective targeting of PDA-FNDs to biomolecules of interest and thus measure thermal conductivity at specific intracellular organelles, for example, inside mitochondria.

The structure of PDA-FND is more uniform (**Fig. 1**) if compared to the other hybrids with similar function, but there is still a significant heterogeneity among particles. Although the Bayesian statistical analysis of the results obtained in different media allows separation of the variance of the temperature rise caused by the inhomogeneity of κ_cell_ from the variance related to the heterogeneity of the particles, removing the latter factor would significantly increase the accuracy with which the value of κ_cell_ could be determined. Moreover, elimination of particle heterogeneity would enable single PDA-FND particle measurement of *κ* at a specific location. We assume that this heterogeneity is related to the intrinsic property of FNDs such as the size, shape, crystal strain, impurity content, and the number of NV^−^ centers. Particles of FND are all different even if they are drawn from the same batch^66,67^. Production of physically and chemically uniform FNDs much needed for applications is yet to be achieved. The problem caused by the heterogeneity of FND could be effectively mitigated, if the same PDA-FND particle were used in different environment. As a proof of principle, a series of ODMR spectra were obtained for four PDA-FNDs attached to the bottom of the flow cell coated by PEI, while the flow cell which has been in air was filled sequentially with water, and then, with oil (**Supplementary Information** and **Fig. S13**). Unfortunately, effective delivery of a preselected particle into a cell is still an unresolved challenge.

In summary, we have successfully measured the intracellular thermal conductivity of single living cells, *κ*_cell_, for the first time. The average value of *κ*_cell_ is an order of magnitude smaller than *κ*_water_ but is close to *κ*_oil_. Further analysis shows variation in *κ*_cell_ due to significant variation of the intracellular media. Our experimental results provide essential information for theoretical considerations of the role of heat transfer in cells. In particular, the hypothesis about heat-mediated intracellular signalling gains additional support and, perhaps, will result in better understanding of control and regulation processes of heat in living organisms at the single-cell level.

## Materials and methods

### Chemicals and materials

Dopamine hydrochloride (H8502-5G) and nocodazole (M1404-2MG) were purchased from Sigma-Aldrich Co. ltd. (St. Louis, MO, USA). TRIS (hydroxymethyl aminomethane) (35434-76) was a product from Nacalai tesque Inc. (Kyoto, Japan). Sodium azide (NaN_3_) (195-11092) was a product from Fujifilm Wako Pure Chemical Corporation (Osaka, Japan). Hoechst 33342 (346-07951) and cell counting kit-8 (CCK-8) (347-07621) were purchased from Dojindo Laboratories (Kumamoto, Japan). Fluorescent nanodiamond (FND, BR100) was purchased from FND Biotech, Inc. (Taipei, Taiwan). LAMP1-mGFP plasmid, which encodes a GFP tagged lysosome marker, lysosomal-associated membrane protein 1 (LAMP1), was a gift from Esteban Dell’Angelica (Addgene plasmid # 34831; http://n2t.net/addgene:34831; RRID: Addgene_34831). Lipofectamine 3000 Transfection Kit (3000-015) was a product from Invitrogen (CA, USA). Dulbecco’s modified eagle medium (DMEM) (11965-092), 0.25% Trypsin-EDTA (25200-056), phosphate buffered saline (PBS) (10010-023), penicillin and streptomycin (15140-122), fetal bovine serum (FBS) (10270-106), Alexa Fluor 488 NHS Ester (A20000), CellMask Green (C37608), ER-tracker (E34251) and Golgi-tracker (B22650) were products from Thermo Fisher Scientific (MA, USA).

### Synthesis and characterization of PDA-FND particles

PDA-FNDs were prepared according to the protocol reported previously with minor modifications^44^. To synthesize PDA-FND nanoparticles, FNDs were mixed with a freshly prepared dopamine hydrochloride solution in 10 mM Tris-HCl buffer, pH 8.5, with varied concentrations of 50 μg/ml, 300 μg/ml and 500 μg/ml, and allowed to incubate for 16 hours under vigorous shaking at 298 K on a thermo-shaker (TS-100C, bioSan Medical-biological Research & Technologies, Riga, Latvia). Then, the resulting PDA-FNDs were collected by centrifugation at × 3000 *g* for 10 minutes at 298 K. Finally, the nanoparticles were dispersed in Milli-Q water for further use. Dynamic light scattering (DLS) and Zeta potential of the nanoparticles were measured with a Malvern Zetasizer instrument (Zetasizer nano ZSP, Malvern Instruments Limited, Worcestershire, UK). H-7650 transmission electron microscope (TEM) (Hitachi high technology, Tokyo, Japan) operating at an accelerating voltage of 80 kV was adopted to characterize the size, PDA shell thickness and structure of FNDs and PDA-FNDs. The TEM samples were prepared by dropping 5 μl of solution onto a 400-mesh carbon film-coated copper grids (24945-50, Polysciences, Inc., PA, USA), which were glow-discharged by a plasma cleaner. The grids were then air-dried and subjected to TEM observation.

### TEM image analysis

Image J was utilized to analyze the TEM images of FND and PDA-FND. The radius of FND was defined as half of the average length of a horizontal line and a vertical line, which were drawn across the FND to pass through its center. To measure the thickness of PDA layer, tangent lines were drawn at the both ends of the vertical and horizontal lines. The length of the four-line segments, which were vertical to the tangent lines across the PDA layer, was measured. The thickness of the PDA layer was determined by averaging the four measurements.

### Cell culture and cytotoxicity assay (CCK-8)

HeLa cells were seeded on 100 mm tissue culture dish (TRP Techno Plastic Products AG, Switzerland) and were cultured in DMEM supplemented with 10% FBS, 100 IU/ml penicillin and 100 μg/ml streptomycin, under a humidified atmosphere containing 5% CO_2_.

To evaluate the cytotoxicity of PDA-FND, HeLa cells were seeded to 96-well plates (Thermo Fisher Scientific) at the density of 5000 per well. After 24 hours, the cells were treated with PDA-FNDs in various concentrations, and incubated for four hours at 310 K. Then the PDA-FNDs were washed out and the samples were subjected to overnight incubation at 310 K. The next day, the cell viability was determined using a micro plate reader (Thermo Scientific Multiskan FC, Thermo Fisher Scientific Oy, Ratstie, Finland) according to the manufacture’s protocol of CCK-8, in which water-soluble tetrazolium salt, WST-8 (2-(2-methoxy-4-nitrophenyl)-3-(4-nitrophenyl)-5-(2,4-disulfophenyl)-2H-tetrazolium), is reduced by dehydrogenase activities in cells to yield a yellow-color formazan dye. The 450 nm absorption before the addition of CCK-8 solution was deducted to subtract the absorption by PDA-FNDs inside the HeLa cells. The viability of HeLa cells treated with PDA-FNDs was expressed as a percentage of the viability of HeLa cells grown in the absence of PDA-FNDs.

### Fluorescent staining of subcellular organelles and confocal fluorescence microscopy

HeLa cells were seeded in 35 mm glass-based dishes (IWAKI, Shizuoka, Japan). PDA-FNDs were dispersed in culture medium and were applied to HeLa cells. After four hours incubation at 310 K, cells were stained with CellMask Green Plasma Membrane Stain (522 nm and 535 nm) (1:2000 dilution) and 1 μg/ml Hoechst 33342 (Ex and Em; 355 nm and 461 nm) to visualize plasma membrane and nucleus, respectively, according to the manufacture’s procedures. Cells were also stained with 500 nM ER-tracker Green (504 nm and 511 nm) for endoplasmic reticulum (ER), with 250 nM BODIPY FL C5-ceramide complexed to BSA (505 nm and 511 nm) for Golgi apparatus, or with 5 μM MitoTracker green FM (490 nm and 516 nm) for mitochondria to examine colocalizations with these intracellular organelles. To inhibit the energy-dependent endocytosis process, HeLa cells were treated with 10 mM NaN_3_ for 30 min, then the cells were incubated with PDA-FNDs for four hours, followed by CellMask Green staining. Then, the prepared specimens were observed under Leica TCS SP8 confocal microscope (Leica Microsystems, Germany) with a 63 × objective lens (HC PL APO CS2 63×/1.40 OIL).

### Fluorescence activated cell sorting (FACS)

HeLa cells were seeded to 96-well plate, and PDA-FNDs were loaded to the cells once they reached confluency. After four hours incubation at 310 K, unbound PDA-FNDs were washed out and the cells were collected by centrifugation at × 140 *g* following Trypsin-EDTA treatment. The cell pellet was washed twice in PBS and resuspended in 100 μl PBS. The samples were then subjected to FACS analysis with the excitation at 561 nm and the emission at 675-715 nm by an Attune NxT Flow Cytometer (Thermo Fisher Scientific). For the NaN_3_ treatment group, HeLa cells were pretreated with 10 mM NaN_3_ for 30 minutes before the loading of PDA-FND, and 10 mM NaN_3_ was present in all the solutions during incubation, sample collection and measurement.

### Overexpression of LAMP1-mGFP in HeLa cells

HeLa cells were transfected with LAMP1-mGFP plasmid by using Lipofectamine 3000 Transfection Kit. After 24 hours, the cells were loaded with 10 μg/ml PDA-FNDs and incubated at 310 K for four hours, then the unbound PDA-FNDs were washed out and the cells were subjected to overnight incubation followed by the fluorescence confocal microscopy.

### Temperature measurement using FNDs through optically detected magnetic resonance microscopy

The temperature was measured using FNDs and a home-built microscope, which was reported previously^33^. Briefly, a continuous Nd:YAG laser at 532 nm illuminated FNDs in PDA-FND nanoparticles to initialize and read out the spin state of NVCs on an inverted microscope system (Nikon, Ti-E). Fluorescence from the NVCs was captured by an oil-immersion objective lens (Nikon SR HP Apo TIRF, 100×, NA 1.49), passed through a long-pass filter to cut off the excitation light, and imaged by an electron-multiplying charge-coupled device camera (EMCCD, Andor, iXon860). A two-turn copper coil with a diameter of approximately 1 mm was placed just above the coverslip to irradiate the sample with microwaves (MWs) at frequencies near the electron-spin resonance in NVCs. A gate pulse provided by a pulser (Berkley Nucleonics Corporation, 565) activated a MW generator (Agilent, E8257D) to output a MW pulse, which was then amplified by linear MW power amplifiers (Mini-Circuits, PAN35-5A and ZHL-16W-43+) and transmitted through a coaxial cable to the MW coil. The pulser and the image acquisition by the EMCCD camera were synchronized by a PC (Dell T3500, XT, USA) with LabVIEW software (National Instruments, TX, USA). ODMR spectra were recorded for a range of MW frequencies as the difference between the fluorescence intensities with and without MW irradiation. The temperature of the sample was set at 300 K by a stage-top incubator (TOKAI HIT, TPi-108RH26) on a motorized stage.

### Evaluation of the photothermal effect of PDA-FNDs

Glass-based dish was first coated by thin layer of polyethylene imine (PEI) in order to improve the adhesion of PDA-FND, and then PDA-FNDs were scattered on the dish. Fluorescence images were recorded while the MW frequency was digitally swept across the range from 2860 to 2978 MHz with an increment of 0.178 MHz. Exposure time of the EMCCD camera was altered from 0.05 s to 0.005 s depending on the laser intensity. A set of images (two for cell experiment and 64 in other cases) were accumulated per MW frequency, and two scans (32 for cell experiment) were performed. Through this process, ODMR spectra were obtained from isolated PDA-FND particles located in the field of view. Each spectrum was fitted to the six-parameter superposition of two Lorentzian functions.

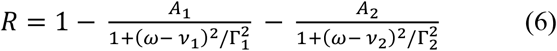

where the MW frequency *ω* assumes a discrete set of values *ω_n_* incremented by the 0.178 MHz. The *χ*^2^-function of six parameters in **Eq. (6)** is defined as

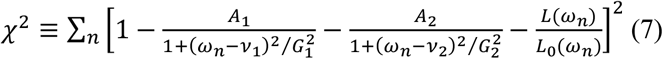

where *L*(*ω_n_*) and *L*_0_(*ω_n_*) are the measured luminescence intensities with MW power “on” and “off”, respectively. The minimum of *χ*^2^ is found numerically for each measured ODMR spectrum. The corresponding values of all parameters are their best posterior estimates. The second derivatives of *χ*^2^ at the minimum are used to estimate the covariance matrix whose diagonal elements are the variances for each parameter in **Eq. (6)** and they define the precision for each fitted parameter (details can be found in elsewhere^68^, for example).

The temperature change in each FND has been determined relative to the temperature at a reference excitation laser power, which was either 7.3 mW or 0.95 mW. For example, d*T*_7.3→*P*_ = *γ*Δ*D*_gs_, where and Δ*D*_gs_ is the change of *D*_gs_ = (*v*_1_ + *v*_2_)/2 [45] when the laser power changes from 7.3 mW to P.

All the numerical simulations were performed using a standard library of MATLAB.

### ODMR measurement of PDA-FNDs inside HeLa cells

HeLa cells were cultured on 35-mm glass-based dish and incubated with 0.5 μg/ml PDA-FNDs for four hours. Then the PDA-FNDs were washed out and the cells were cultured at 310 K overnight. Next day the cells were washed once with serum-free DMEM, and the medium was replaced by 2 ml of 10 μM nocodazole dissolved in phenol red-free DMEM. After 30-min. incubation at 310 K, the cells were subjected to ODMR measurement.

### Measurement of the thickness of PEI layer using high-speed AFM combined with fluorescence microscopy

PEI solution containing 10 μg/ml of Alexa Fluor 488 NHS Ester (Alexa488-PEI) was applied on a glass coverslip to prepare PEI layer. First, the coverslip was observed under the fluorescence microscope to identify the edges of the Alexa488-PEI layer. A part of the coverslip containing the edge of Alexa488-PEI layer was cut into a fragment of approximately 2-mm square. The fragment was glued onto the AFM sample stage. Then, measurements of the thickness of the Alexa488-PEI layer were performed with a laboratory-built high-speed AFM (HS-AFM) apparatus. The HS-AFM is similar to the HS-AFM previously reported^69,70^, whereas the current apparatus was customized to capture a fluorescence image of the sample on the AFM stage. The HS-AFM was equipped with small cantilevers (k = 0.1 – 0.2 N/m, f = 800 – 1200 kHz in water (Olympus)) and operated in tapping mode. The cantilever was approached onto the edge of the Alexa488-PEI layer with the guidance of the fluorescence image of the AFM stage. HS-AFM observation was performed under ultrapure water at the room temperature. The height of the Alexa488-PEI layer in AFM images was analyzed using Igor Pro (WaveMetrics).

## Acknowledgements

We appreciate Dr. Yohsuke Yoshinari for setting up the ODMR microscope, and Dr. Fuminori Sugihara for his technical advice on FACS. We also thank Prof. Shin’ichi Ishiwata for valuable comments and discussion. This work was supported by Grant-in-Aid for JSPS Research Fellow A18J002870 (to SS), Early-Career Scientists 19K16089 (to SS), and by ATI Research Grant RG3004 (to SS), by JSPS KAKENHI JP15H05931, JP18H01838 (both to YH), JP15K05251 and JP16KK0105 (both to MS), by the Japan Science and Technology Agency JPMJPR15F5 (to MS) and JPMJPR15FD (to HY), by the Human Frontier Science Program RGP0047/2018 (to TP and MS), by Kurita Water and Environment Foundation 17D002 (to MS), by Iketani Science and Technology Foundation 0301009-A (to MS).

## Author contributions

S.S., C.Z., J.C.Y.K., T.P., Y.H. and M.S. designed the experiments. S.S. and C.Z. synthesized and characterized PDA-FND. C.Z. carried out the cell viability tests and intracellular localization analysis. S.S. carried out the ODMR measurements. H.Y. carried out the HS-AFM measurements. T.P. carried out the theoretical studies and numerical simulations. S.S., C.Z., H.Y., T.P. and M.S. analyzed the data. S.S., C.Z., H.Y., T.P. Y.H. and M.S. wrote the paper with input from J.C.Y.K.

