## Supplementary Information for "*In situ* measurement of intracellular thermal conductivity using heater-thermometer hybrid diamond nanosensor"

### Analytical model

A PDA-FND complex and surrounding media with a spherical symmetry is shown in **Fig. 4A**. The macroscopic differential equation for the steady-state temperature $T(r)$ reads in this case

$$-\kappa\frac{dT}{dr}=\frac{\dot{Q}(r)}{4\pi r^2} \tag{S1}$$

where $\kappa$ is the thermal conductivity. The heat power $\dot{Q}(r)=\epsilon IV$, where $I$ is the laser irradiance at the location of the particle, $\epsilon$ is the absorption coefficient of PDA, and $V$ is the volume of PDA enclosed by radius $r$ (dashed line in **Fig. 4A**). One can integrate **Eq. (S1)** to obtain $\Delta T$, the change of temperature in the FND caused by the change of irradiance $\Delta I$.

Integration of **Eq. (S1)** results in

$$\Delta T=\frac{\epsilon\Delta I}{3}\left(\int_{r_\mathrm{i}}^{\infty}\frac{r_\mathrm{p}^3-r_\mathrm{c}^3}{\kappa_\mathrm{h}r^2}\mathrm{d}r+\int_{r_\mathrm{p}}^{r_\mathrm{i}}\frac{r_\mathrm{p}^3-r_\mathrm{c}^3}{\kappa_\mathrm{i}r^2}\mathrm{d}r+\int_{r_\mathrm{c}}^{r_\mathrm{p}}\frac{r^3-r_\mathrm{c}^3}{\kappa_\mathrm{p}r^2}\mathrm{d}r\right)$$

$$=\frac{\epsilon\Delta I}{3}\left(\frac{r_\mathrm{p}^3-r_\mathrm{c}^3}{r_\mathrm{i}\kappa_\mathrm{h}}+\left[\frac{r_\mathrm{p}^3-r_\mathrm{c}^3}{r_\mathrm{p}\kappa_\mathrm{i}}-\frac{r_\mathrm{p}^3-r_\mathrm{c}^3}{r_\mathrm{i}\kappa_\mathrm{i}}\right]+\left[\frac{r_\mathrm{p}^2-r_\mathrm{c}^2}{2\kappa_\mathrm{p}}+\frac{r_\mathrm{c}^3}{r_\mathrm{p}\kappa_\mathrm{p}}-\frac{r_\mathrm{c}^2}{\kappa_\mathrm{p}}\right]\right)$$

$$=\frac{\epsilon\Delta I}{3}\left(\frac{r_\mathrm{p}^3-r_\mathrm{c}^3}{r_\mathrm{i}\kappa_\mathrm{h}}+\frac{\left(r_\mathrm{p}^3-r_\mathrm{c}^3\right)\left(r_\mathrm{i}-r_\mathrm{p}\right)}{r_\mathrm{i}r_\mathrm{p}\kappa_\mathrm{i}}+\frac{r_\mathrm{p}^3-3r_\mathrm{c}^2r_\mathrm{p}+2r_\mathrm{c}^3}{2r_\mathrm{p}\kappa_\mathrm{p}}\right)$$

$$=\frac{\epsilon\Delta I}{3}\left(\frac{r_\mathrm{p}^3-r_\mathrm{c}^3}{r_\mathrm{p}k\kappa_\mathrm{h}}+\frac{r_\mathrm{p}^3-r_\mathrm{c}^3}{r_\mathrm{p}\kappa_\mathrm{h}}\times\frac{\left(r_\mathrm{i}-r_\mathrm{p}\right)\left(\kappa_\mathrm{h}-\kappa_\mathrm{i}\right)}{r_\mathrm{i}\kappa_\mathrm{i}}+\frac{r_\mathrm{p}^3-3r_\mathrm{c}^2r_\mathrm{p}+2r_\mathrm{c}^3}{2r_\mathrm{p}\kappa_\mathrm{p}}\right)$$

$$=\Delta T_\mathrm{h}+\Delta T_\mathrm{h}\frac{r_\mathrm{i}-r_\mathrm{p}}{r_\mathrm{i}}\frac{\kappa_\mathrm{h}-\kappa_\mathrm{i}}{\kappa_\mathrm{i}}+\Delta T_\mathrm{p} \tag{S2}$$

where

$$\Delta T_\mathrm{h}=\frac{\epsilon\Delta I}{3\kappa_\mathrm{h}}\times\left(r_\mathrm{p}^2-r_\mathrm{c}^3r_\mathrm{p}^{-1}\right) \tag{S3}$$

depends on the thermal conductivity of the media, but

$$\Delta T_\mathrm{p}=\frac{\epsilon\Delta I}{3\kappa_\mathrm{p}}\times\frac{r_\mathrm{p}^2-3r_\mathrm{c}^2+2r_\mathrm{c}^3r_\mathrm{p}^{-1}}{2} \tag{S4}$$

is independent of the surroundings.

In agreement with experiments (**Fig. 1C and D**), it is assumed that $r_\mathrm{p}=r_\mathrm{c}+\delta r$, where $r_\mathrm{c}\approx\delta r\approx 50$ nm. In this case, the second fraction in **Eq.(S4)** is about three times smaller than $r_\mathrm{p}^2-r_\mathrm{c}^3r_\mathrm{p}^{-1}$ in **Eq. (S3)**, and hence $\Delta T_\mathrm{p}\ll\Delta T_\mathrm{h}$ in air and oil whereas it is not in water where

$\kappa_\mathrm{h} \approx 3\kappa_\mathrm{p}$ (see **Table 1**). The significance of the uncertainty of the particle size is proportional to derivatives of the factors $r_\mathrm{p}^2 - r_\mathrm{c}^3 r_\mathrm{p}^{-1}$ in **Eq. (S3)** and $r_\mathrm{p}^2 - 3r_\mathrm{c}^2 + 2r_\mathrm{c}^3 r_\mathrm{p}^{-1}$ in **Eq. (S4)** over $r_\mathrm{c}$. The derivative of the factor in **Eq. (S4)** is 10-fold smaller than that in **Eq. (S3)** (**Fig. S8**). Therefore, the contribution of $\Delta T_\mathrm{p}$ to the variance of $\Delta T$ is negligible in all investigated media except for water where $\Delta T_\mathrm{p}$ makes an appreciable addition to the variance. If additionally, $\langle \kappa_\mathrm{h}/\kappa_\mathrm{i} \rangle = 1$ and fluctuations of $r_\mathrm{c}$, $r_\mathrm{i}$, and $\kappa_\mathrm{i}$ are uncorrelated, then

$$\sigma_{\Delta T}^2 \approx \langle \Delta T_\mathrm{h} \rangle^2 \frac{\sigma^2_{r_\mathrm{p}^2 - r_\mathrm{c}^3 r_\mathrm{p}^{-1}}}{\langle r_\mathrm{p}^2 - r_\mathrm{c}^3 r_\mathrm{p}^{-1} \rangle^2} + \langle {\Delta T_\mathrm{h}}^2 \rangle \left\langle \left(1 - r_\mathrm{p}/r_\mathrm{i}\right)^2 \right\rangle \sigma^2_{\kappa_\mathrm{h}/\kappa_\mathrm{i}} \tag{S5}$$

This is because

$$\sigma_{x+xyz}^2 = \langle (x + xyz)^2 \rangle - \langle x + xyz \rangle^2 = \langle x^2 \rangle - \langle x \rangle^2 + \langle x^2 y^2 z^2 \rangle = \sigma_x^2 + \langle x^2 \rangle \langle y^2 \rangle \sigma_z^2 \tag{S6}$$

where $x \equiv \Delta T_\mathrm{h}$, $y \equiv 1 - r_\mathrm{p}/r_\mathrm{i}$ and $z \equiv \kappa_\mathrm{h}/\kappa_\mathrm{i} - 1$ are uncorrelated and $\langle z \rangle = 0$.

The first term in **Eq. (S5)** is referenced as $\sigma^2_{\Delta T_\mathrm{h}}$. It is a product of $\langle \Delta T_\mathrm{h} \rangle^2$ and a quantity related to the statistics of PDA-FND nanoparticles. As it is independent of the surroundings, $\sigma^2_{\Delta T_\mathrm{h}} = \langle \Delta T \rangle^2_\mathrm{air} / \langle \Delta T_\mathrm{h} \rangle^2 \times \sigma^2_{\Delta T,\mathrm{air}}$ in all media. The second term in **Eq. (S5)** is $\sigma^2_\mathrm{env}$. By means of the ratios $\kappa_\mathrm{h}/\kappa_\mathrm{i}$ and $r_\mathrm{p}/r_\mathrm{i}$, it depends on fluctuations of the local environment. It is reasonable to assume that $\sigma^2_\mathrm{env} = 0$ in air and other homogeneous media. If this assumption holds, then $\sigma^2_{\Delta T_\mathrm{h},\mathrm{air}} = \sigma^2_{\Delta T,\mathrm{air}}$. If this is not the case, the variation can be quantitatively characterized by the variance of $\kappa_\mathrm{h}/\kappa_\mathrm{i}$, that is by $\sigma^2_{\kappa_\mathrm{h}/\kappa_\mathrm{i}}$. According to the definition of $\sigma^2_\mathrm{env}$ (**Eq. S5**) and the equality $\langle {\Delta T_\mathrm{h}}^2 \rangle = \langle \Delta T_\mathrm{h} \rangle^2 + \sigma^2_{\Delta T_\mathrm{h}}$ one gets

$$\sigma^2_{\kappa_\mathrm{h}/\kappa_\mathrm{i}} = \frac{\sigma^2_\mathrm{env}}{\left(\langle \Delta T_\mathrm{h} \rangle^2 + \sigma^2_{\Delta T_\mathrm{h}}\right)\left\langle \left(1 - r_\mathrm{p} {r_\mathrm{i}}^{-1}\right)^2 \right\rangle} \tag{S7}$$

### Bayesian inferences

Each particle in this experiment has a physical temperature rise of $\Delta T$ which is different from the measured value $\Delta T_\mathrm{m}$ because of the measurement error. It is assumed that the errors have a normal probability distribution with zero mean and standard deviation of $\sigma_\mathrm{m}$. Second, the values of *physical* temperatures $\Delta T$ are different for different particles. This difference can arise from the difference in size of the PDA-FND particles, the thickness of PDA, and from the hypothetical variation of the local thermal conductivity. It is assumed that the variation of $\Delta T$ is also normal and has a mean $\langle \Delta T \rangle$ and a standard deviation $\sigma_{\Delta T}$. We use Bayesian approach to analyze the data ($\Delta T_\mathrm{m}$ and $\sigma_\mathrm{m}$) to infer information about $\langle \Delta T \rangle$ and $\sigma_{\Delta T}$.

According to the Bayes' theorem, the inferred joint probability for $\langle\Delta T\rangle$ and $\sigma_{\Delta T}$ given *data* (*D*) reads

$$P(\langle\Delta T\rangle,\sigma_{\Delta T}^2|D) \propto P(\sigma_{\Delta T}^2)\prod_{m=1}^{N}\int_{-\infty}^{\infty}\exp\left(-\frac{(\Delta T-\Delta T_m)^2}{2\sigma_m^2}\right)\exp\left(-\frac{(\Delta T-\langle\Delta T\rangle)^2}{2\sigma_{\Delta T}^2}\right)\mathrm{d}\Delta T \quad \text{(S8)}$$

The integral in **Eq. (S8)** equals the probability $P(\Delta T_m|\langle\Delta T\rangle,\sigma_{\Delta T},\sigma_m)$. The *prior* distribution of $\langle\Delta T\rangle$ is assumed uniform but we have explicitly included *prior* $P(\sigma_{\Delta T}^2)$. The integration is straightforward and the result reads

$$P(\langle\Delta T\rangle,\sigma_{\Delta T}^2|D) \propto P(\sigma_{\Delta T}^2)\prod_{m=1}^{N}\frac{\exp\left(-\frac{[\langle\Delta T\rangle-\Delta T_m]^2}{2[\sigma_m^2+\sigma_{\Delta T}^2]}\right)}{(\sigma_m^2+\sigma_{\Delta T}^2)^{1/2}} \quad \text{(S9)}$$

With some algebra, this can be transformed in the following expression

$$P(\langle\Delta T\rangle,\sigma_{\Delta T}^2|D) \propto P(\sigma_{\Delta T}^2)\frac{\exp\left(-\frac{\Upsilon_\mathrm{w}+[\langle\Delta T\rangle-T_\mathrm{w}]^2}{2s_\mathrm{w}^2}\right)}{\prod_{m=1}^{N}(\sigma_m^2+\sigma_{\Delta T}^2)^{\frac{1}{2}}} \quad \text{(S10)}$$

where

$$\frac{1}{s_\mathrm{w}^2}\equiv\sum_{m=1}^{N}\frac{1}{\sigma_m^2+\sigma_{\Delta T}^2};\ T_\mathrm{w}\equiv s_\mathrm{w}^2\sum_{m=1}^{N}\frac{\Delta T_m}{\sigma_m^2+\sigma_{\Delta T}^2};\ \Upsilon_\mathrm{w}\equiv s_\mathrm{w}^2\sum_{m=1}^{N}\frac{\Delta T_m^2}{\sigma_m^2+\sigma_{\Delta T}^2}-T_\mathrm{w}^2 \quad \text{(S11)}$$

One can obtain $P(\langle\Delta T\rangle|D)$ and $P(\sigma_{\Delta T}^2|D)$ by integration of $P(\langle\Delta T\rangle,\sigma_{\Delta T}^2|D)$ over $\sigma_{\Delta T}^2$ and $\langle\Delta T\rangle$, respectively (called "marginalization"). The first integral can be done numerically (**Fig. S9**), whereas the integration over $\langle\Delta T\rangle$ is simple and results in the expression

$$P(\sigma_{\Delta T}^2|D) \propto P(\sigma_{\Delta T}^2)\rho_{\sigma_{\Delta T}^2}(\sigma_{\Delta T}^2) \quad \text{(S12)}$$

The function $\rho_{\sigma_{\Delta T}^2}(\sigma_{\Delta T}^2)$

$$\rho_{\sigma_{\Delta T}^2}(\sigma_{\Delta T}^2)\equiv\frac{s_\mathrm{w}\exp\left(-\frac{\Upsilon_\mathrm{w}}{2s_\mathrm{w}^2}\right)}{\prod_{m=1}^{N}(\sigma_m^2+\sigma_{\Delta T}^2)^{1/2}} \quad \text{(S13)}$$

is introduced for convenience.

The case when $\sigma_{\Delta T}^2=\sigma_{\Delta T_\mathrm{h}}^2+\sigma_\mathrm{env}^2$ and the prior $P(\sigma_\mathrm{env}^2,\sigma_{\Delta T_\mathrm{h}}^2)=P(\sigma_\mathrm{env}^2)P(\sigma_{\Delta T_\mathrm{h}}^2)$ is of special interest here. The equation analogous to **Eq. (S10)** reads in this case

$$P(\sigma_{\mathrm{env}}^2, \sigma_{\Delta T}^2|D) \propto P(\sigma_{\mathrm{env}}^2) P(\sigma_{\Delta T_{\mathrm{h}}}^2) \rho_{\sigma_{\Delta T}^2}(\sigma_{\Delta T_{\mathrm{h}}}^2 + \sigma_{\mathrm{env}}^2) \quad \text{(S14)}$$

The marginalized probability $P(\sigma_{\mathrm{env}}^2|D)$ is expressed by the integral over $\sigma_{\Delta T_h}^2$

$$P(\sigma_{\mathrm{env}}^2|D) \propto P(\sigma_{\mathrm{env}}^2) \int P(\sigma_{\Delta T_{\mathrm{h}}}^2) \rho_{\sigma_{\Delta T}^2}(\sigma_{\Delta T_{\mathrm{h}}}^2 + \sigma_{\mathrm{env}}^2) d\sigma_{\Delta T_{\mathrm{h}}}^2 \quad \text{(S15)}$$

Remarkably, $P(\sigma_{\mathrm{env}}^2|D)$ is a mathematical cross-correlation function.

**Equation (S15)** is used to determine $P(\sigma_{\mathrm{env}}^2|D)$ in water, oil and cells. The functions $\rho_{\sigma_{\Delta T}^2}(x)$ are calculated using Eq. (S13) and are shown in **Fig. S10** while $P(\sigma_{\Delta T_{\mathrm{h}}}^2)$ is obtained using the relation $\sigma_{\Delta T_{\mathrm{h}}}^2 = \langle \Delta T \rangle_{\mathrm{air}}^2 / \langle \Delta T_{\mathrm{h}} \rangle^2 \times \sigma_{\Delta T,\mathrm{air}}^2$ and the distribution $P(\sigma_{\Delta T_{\mathrm{h}}}^2)$ for air (calculated using **Eq. (S12)**). Distributions $P(\sigma_{\mathrm{env}}^2|D)$ obtained in such a way are shown in **Fig. 4D**.

### Sequential measurement for directly comparing the temperature increase with a fixed rate of heat flow using single PDA-FNDs in air, water, and oil.

Among the 20-30 measurements at each condition shown in **Fig 4B**, there were values significantly deviated from the mean (for example in oil) which may have been caused by undesirable factors such as aggregation or heterogeneity among particles. On the other hand, the issue caused by the intrinsic irregularity to the particles can be ignored without relying on a large number of measurements, if we can evaluate different environment using the same FND particles. Taking advantage of the photostability of PDA-FND showing no photobleaching (**Figs. 2C, S5**), we finally demonstrated a sequential measurement using the same particles (**Fig. S13**). A series of ODMR spectra were obtained for four PDA-FNDs attached to the bottom of the flow cell coated by PEI, while the flow cell which has been in air was filled sequentially with water, and then, with oil. The method will allow us to ignore the unfavorable intrinsic heterogeneity among FND particles, and thus can provide a simple, facile, and reliable measurement. In air, temperature rise at laser intensity of 25 mW in reference to the excitation at 7.3 mW, $\mathrm{d}T_{7.3\rightarrow 25}$, was approximately 4-7 K. The temperature rise in water was significantly reduced to ca. 1 K while those in oil was partially recovered to give ca. 2 K. These temperature rises in each medium are in good accordance with the data in **Fig 4B**, which implies that respective thermal conductivities ($\kappa$) of air, water, and oil (**Table 1**) are reproducible in single PDA-FNDs.

**Supplementary Figures**

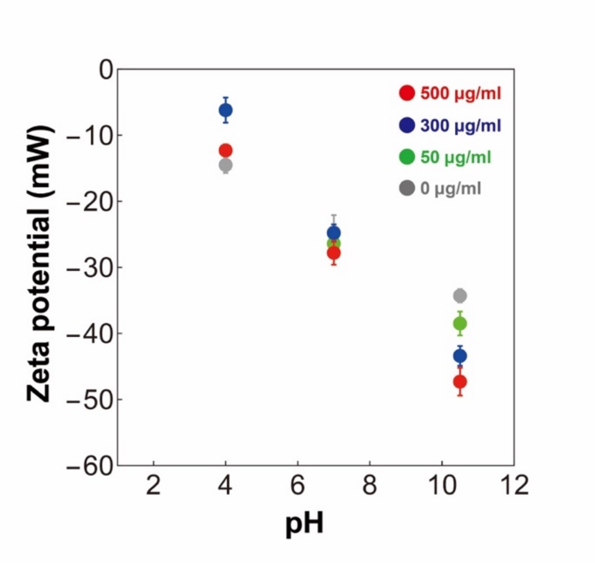


**Figure S1. Zeta potentials of FNDs (grey) and PDA-FNDs prepared with 50 (green), 300 (blue), and 500 µg/ml (red) of dopamine hydrochloride solutions measured at pH 4.0, pH 7.0 and pH 10.5.**

Each five preparation of PDA-FND was subjected for the measurements at pH 4.0, pH 7.0 and pH 10.5. For FND, measurements were repeated for four times. Plots and bars respectively indicate the means and the standard deviations.

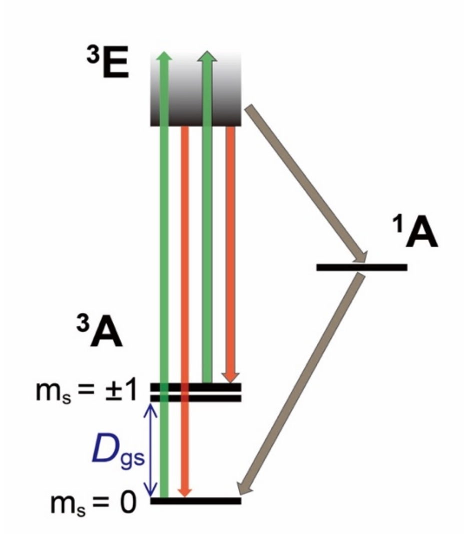


**Figure S2. Energy diagram of the $NV^-$ spin sublevels in nanodiamonds.**

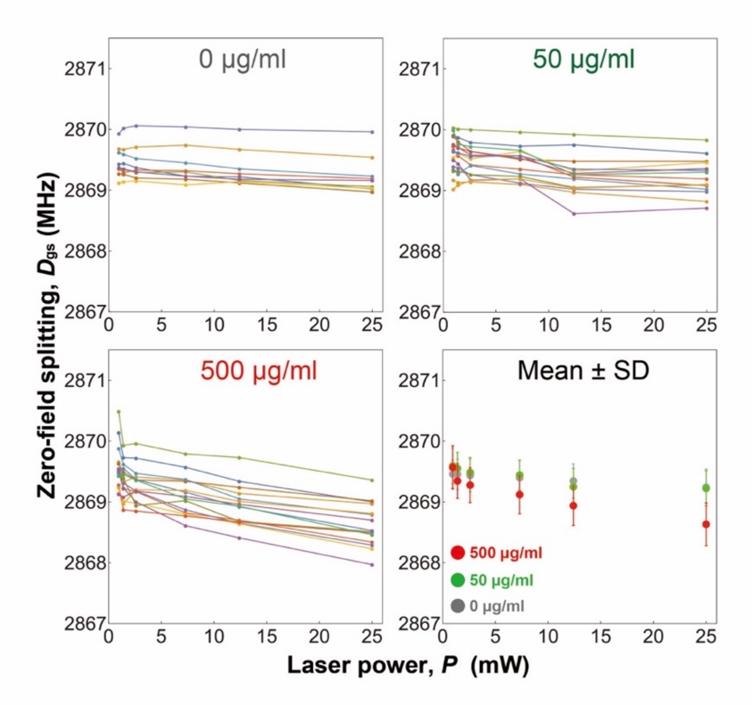


**Figure S3. Correlation between the zero-field splitting of NVC in FND, $D_{gs}$, and the laser power.**

The values of $D_{gs}$ in individual PDA-FNDs prepared with three different concentrations of dopamine hydrochloride solutions as indicated at the top of each panel, and the mean values. Colored plots and connecting lines that are shown in the same color code were obtained sequentially from individual spots in fluorescence images.

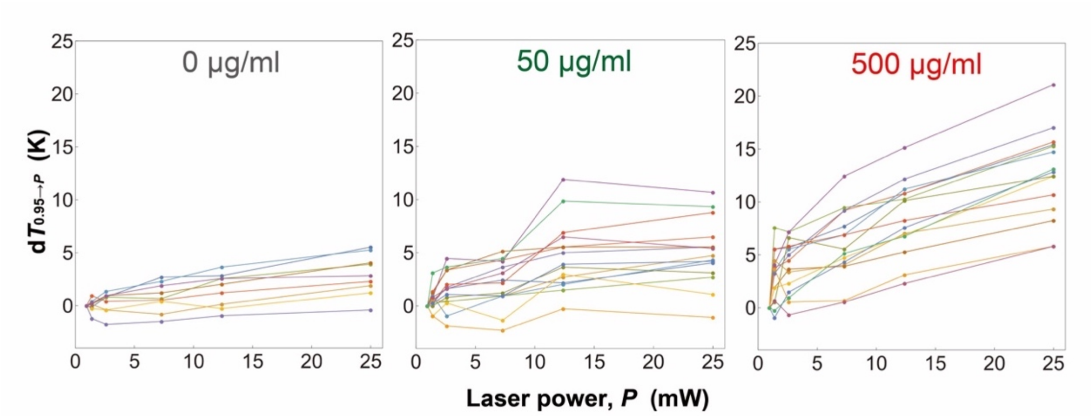


**Figure S4. Correlation between $dT_{0.95 \to P}$ determined by the $D_{gs}$ values in Fig. S3 and the laser power for PDA-FNDs prepared with 0, 50, and 500 µg/ml of dopamine hydrochloride solutions.**

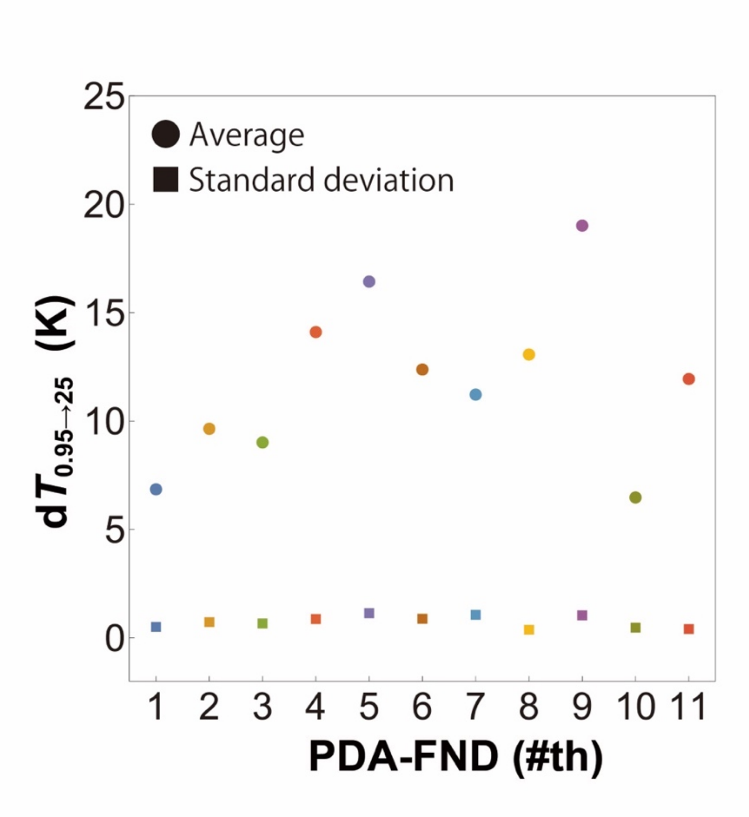


**Figure S5. The means (squares) and the standard deviations (rectangles) of d$T_{0.95\rightarrow25}$ in 11 individual PDA-FNDs for nine times of sequential measurements at 25 mW shown in Fig. 2C in the main text.** Data in **Fig. 2C** was analyzed and then plotted with the same color code.

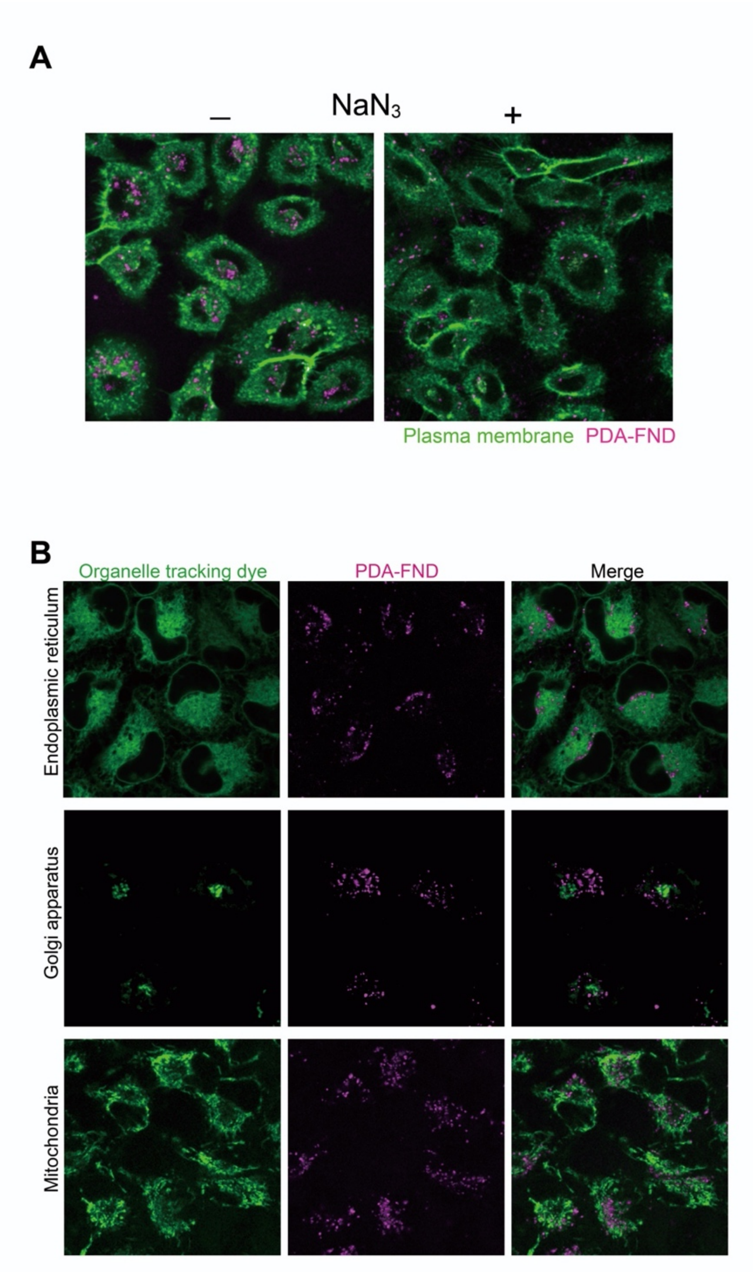


**Figure S6. Internalized PDA-FNDs imaged with various markers of subcellular organelles in HeLa cells**

(A) Confocal microscopy images showing single optical sections of HeLa cells incubated with PDA-FNDs for four hours in the presence (right) and absence (left) of 10 mM $NaN_3$. Green, CellMask Green staining for HeLa cell plasma membrane. Magenta, PDA-FND. Dimension of images, 184.5 μm × 184.5 μm. (B) Single slices of confocal fluorescence microscopy images of PDA-FNDs, tracking dyes for endoplasmic reticulum, Golgi apparatus or mitochondria, and their merges. Dimension of images; 92.3 μm × 92.3 μm. No clear colocalization with endoplasmic reticulum (ER), Golgi apparatus, or mitochondria was observed.

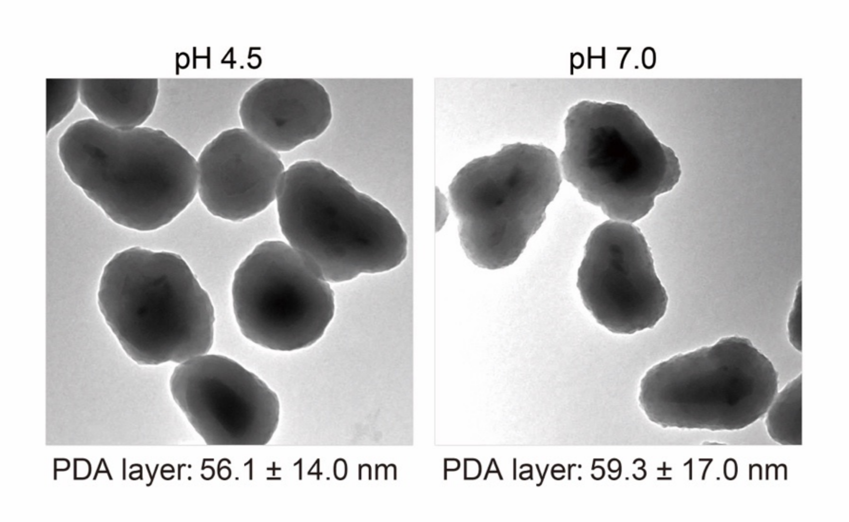


**Figure S7. TEM image analyses of PDA-FNDs incubated overnight at pH4.5 or pH7.0.** Representative TEM images (874 × 874 nm). The thicknesses of PDA layers were measured from the TEM images of PDA-FNDs that have been incubated for overnight at pH 4.5 (n = 76 particles) or pH 7.0 (n = 83) and are presented below the respective images. No significant difference between the thickness of PDA layers in these two groups have been observed, which implies that the PDA layer of PDA-FND was preserved in lysosomes.

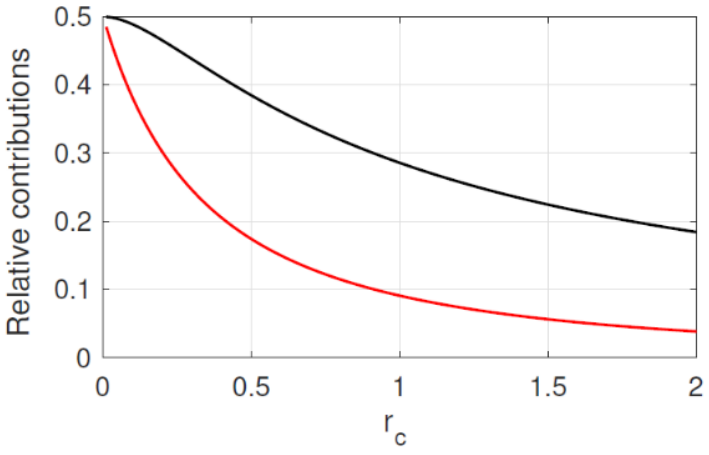


**Figure S8. Relative contributions of $\Delta T_h$ and $\Delta T_p$.**

Black and red lines are $\left(r_p^2 - 3r_c^2 + 2r_c^3 r_p^{-1}\right)/2$ divided by $r_p^2 - r_c^3 r_p^{-1}$, and the ratio of derivatives of this expression over $r_c$. The unit of length is $r_p - r_c$.

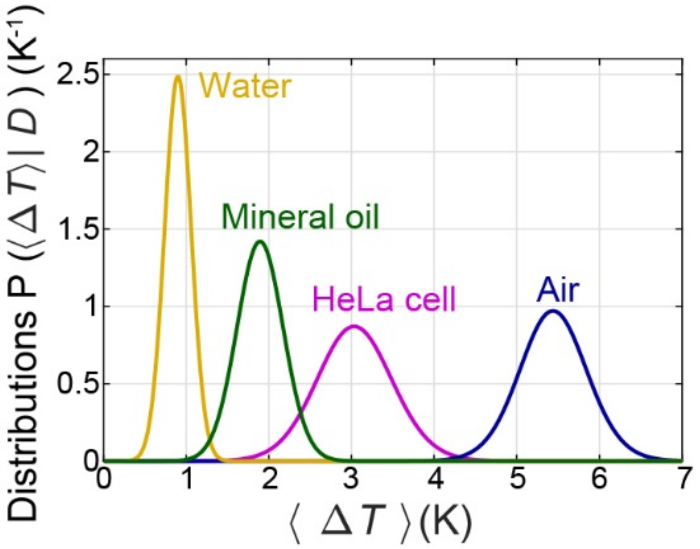


**Figure S9. Probability distribution $P(\langle\Delta T\rangle|D)$ inferred from the results displayed in Fig. 4B.**

The curves from left to right represent water (yellow), mineral oil (green), HeLa cell (magenta), and air (blue).

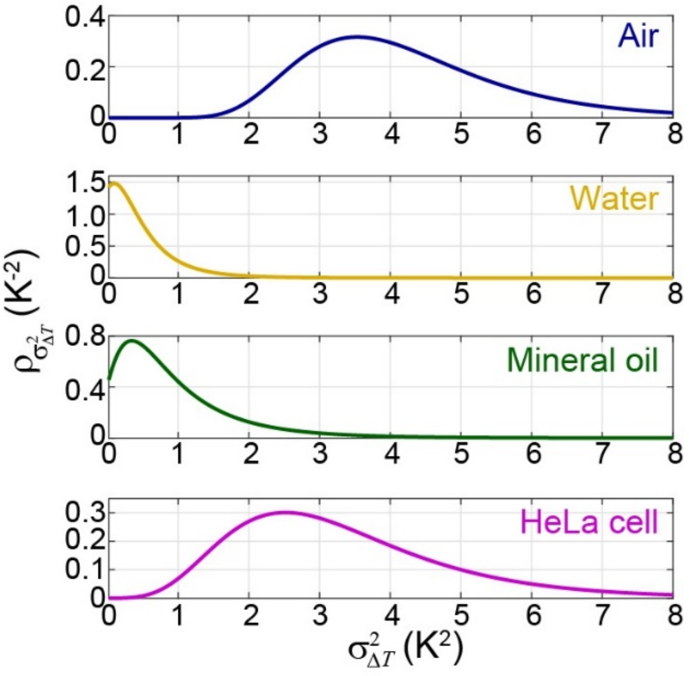


**Figure S10. Posterior probability distributions $\rho_{\sigma^2_{\Delta T}}(\sigma^2_{\Delta T})$**

Posterior probability distributions $\rho_{\sigma^2_{\Delta T}}(\sigma^2_{\Delta T})$ for air, water, oil and cells (from top to bottom) deduced using **Eq. (S13)** and the data in **Fig. 4B**.

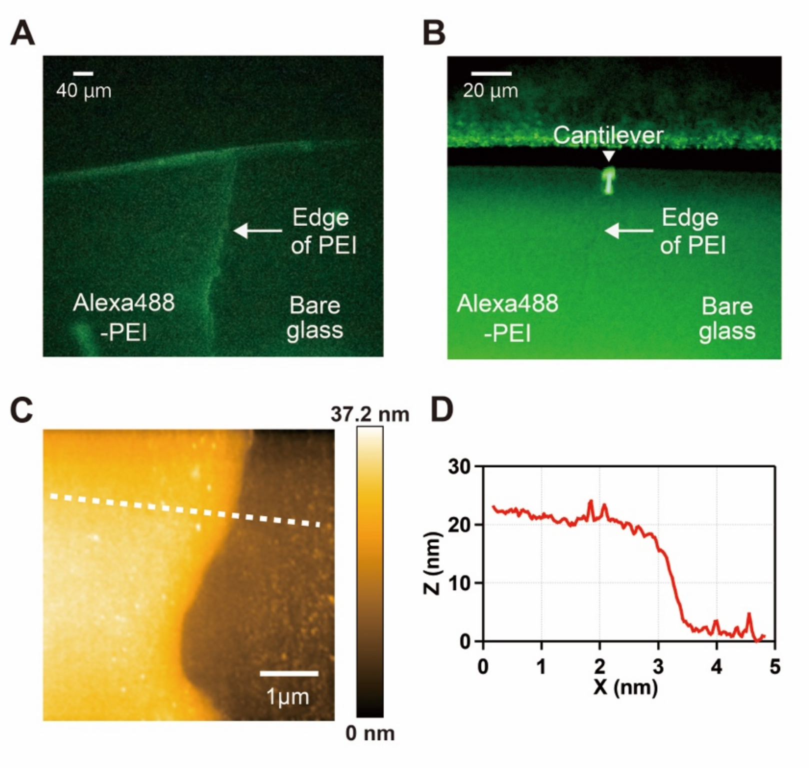


**Figure S11. Measurement of PEI thickness using high-speed AFM (HS-AFM) combined with fluorescence microscopy in aqueous condition**

(A) Fluorescence image of Alexa488-PEI layer coated over a glass coverslip. The edge of Alexa488-PEI layer is visible via the Alexa488 fluorescence. (B) Fluorescence image of the Alexa488-PEI layer and a HS-AFM cantilever. The field of view that is nearly the same as that of (*A*) was captured by another fluorescence microscope which is equipped with the HS-AFM. The edge of Alexa488-PEI layer is visible in the middle of the image. Rectangular shaped AFM cantilever is positioned just above the edge of Alexa488-PEI layer. (C) HS-AFM image of the PEI edge. The scan area was 5 µm × 5 µm. (D) Cross-sectional profile along the dotted line indicated in (*C*).

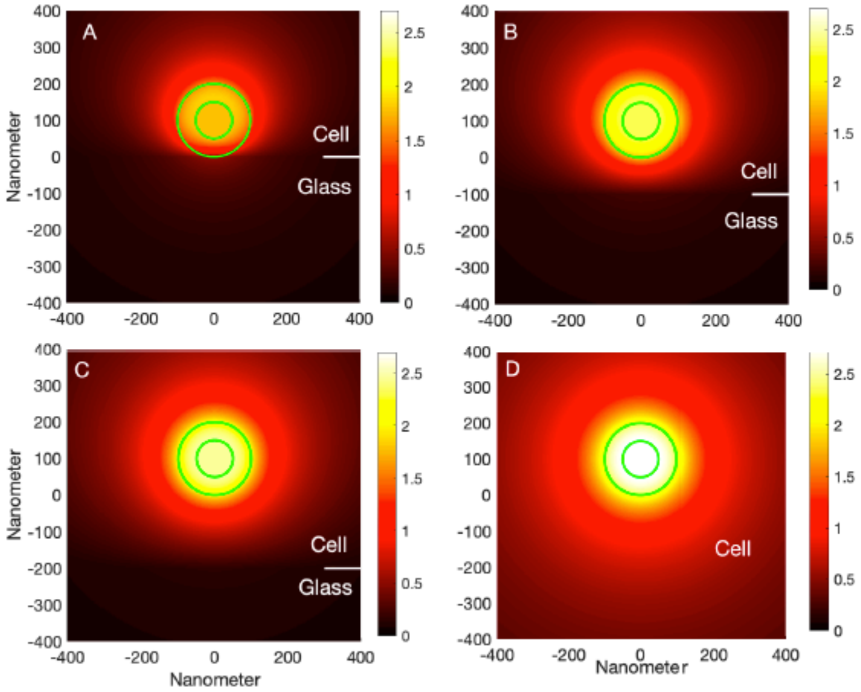


**Figure S12. Numerical simulations of the temperature distributions in and around a PDA-FND particle in a cell on a glass substrate.**

From A to D: The gap between the cell and the glass is set as 0, 0.1, 0.2 and 4 microns and the corresponding temperatures of the FND are 1.8 K, 2.3 K, 2.5 K, and 2.7 K, respectively. The thermal conductivity $\kappa_{cell}$ = 0.12 $Wm^{-1}K^{-1}$. The white horizontal lines in A – C show the interface between the cell and the glass in cases A, B, and C.

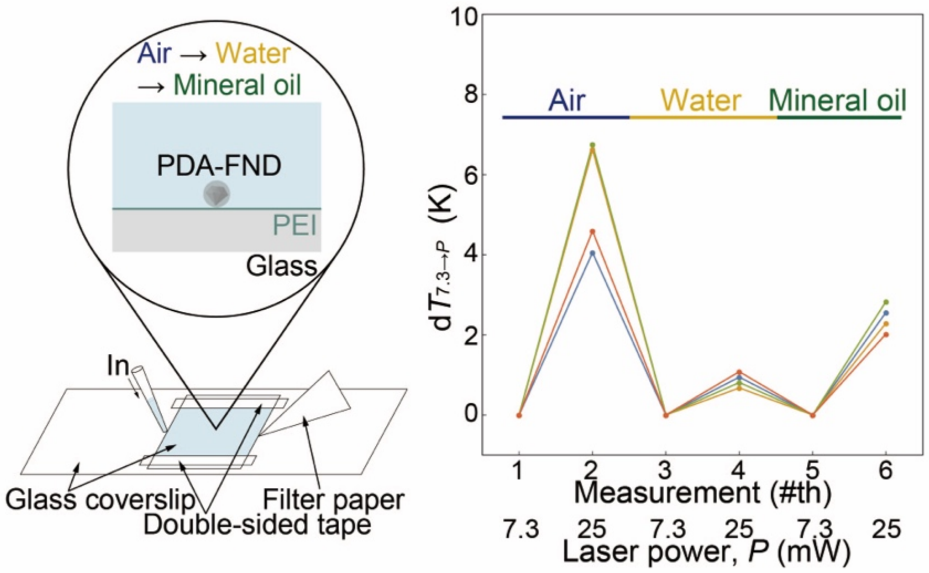


**Figure S13**. **Sequential measurement of $\mathbf{d}T_{7.3\to P}$ using single FND particles.**

Left, schematic illustration of the flow cell where ODMR measurements were performed in air, water, and oil conditions in a successive manner. Right, typical example of successive ODMR measurements of temperature rises in individual PDA-FNDs (n = 4) at different laser powers in air, water and oil. Temperature rise at the laser power of *P* mW, is plotted with reference to the value at 7.3 mW in each medium. Plots that are shown and connected by lines in a same color code represent values obtained from each PDA-FND.

**Supplementary Tables**

**Table S1**. Dynamic light scattering (DLS) analysis of different preparations of FNDs and PDA-FNDs prepared with 500 μg/ml of dopamine hydrochloride solution

| Preparations | FND | | PDA-FND | |
|---|---|---|---|---|
| | Number mean (nm) | SD* (nm) | Number mean (nm) | SD* (nm) |
| 1 | 107 | 29 | 193 | 48 |
| 2 | 105 | 29 | 182 | 43 |
| 3 | 97 | 29 | 190 | 47 |
| 4 | 108 | 28 | 163 | 34 |
| 5 | 107 | 28 | 184 | 44 |
| Mean | 105 | - | 182 | - |

SD*; standard deviation

**Table S2**. Experimental results and numerical simulations of $\langle \Delta T \rangle$ in air, water and oil. Values of $\kappa$ are the literature values. $\kappa_{PDA}$ = 0.2 $Wm^{-1}K^{-1}$. $\dot{q}$ = 100 μW $\mu m^{-3}$ or $\dot{q}$ = 106 μW $\mu m^{-3}$ (in the square brackets)

| | $\langle \Delta T \rangle$ (experiment) K | $\langle \Delta T \rangle$ (simulation) K | $\langle \Delta T \rangle$ (simulation)* K |
|---|---|---|---|
| Air | 5.4 ± 0.8 | 5.49 [5.82] | 5.17 [5.48] |
| Water | 0.9 ± 0.3 | 0.87 [0.92] | 0.83 [0.88] |
| Oil | 1.9 ± 0.6 | 1.81 [1.92] | 1.70 [1.80] |

*Obtained without a 20-nm-thick layer of PEI ($\kappa_{PEI}$ = 0.2 $Wm^{-1}K^{-1}$) on the top of the glass substrate; i.e. between the glass and the PDA-FND.
